# Role of T cells in intrauterine administration of activated peripheral blood mononuclear cells in recurrent implantation failure

**DOI:** 10.1101/2021.01.06.425452

**Authors:** Guillaume Ricaud, Cathy Vaillancourt, Véronique Blais, Marjorie Disdier, Fabien Joao, Bruno Johnson, Moncef Benkhalifa, Pierre Miron, Jacques Bernier

**Affiliations:** Institut National de la Recherche Scientifique (INRS)-Centre Armand-Frappier Santé Biotechnologie, 531 boul. Des Prairies, Laval, Québec H7V 1B7, Canada; Centre d’aide médicale à la procréation FERTILYS, 1950 rue Maurice-Gauvin, Laval, Québec, H7S 1Z5, Canada; Université Picardie Jules Verne, Médecine et Biologie de la Reproduction et laboratoire PERITOX, CBH-CHU Amiens Picardie, 1 Rond-Point du Professeur Christian Cabrol, Amiens, France, 80054

**Keywords:** Endometrium immunomodulation, *In vitro* fertilization, Repeated implantation failure, Peripheral blood mononuclear cells, Th cells

## Abstract

Over the last few years, intrauterine administration of autologous peripheral blood mononuclear cells (PBMC) has been proposed as new immunotherapy for patients with unexplained recurrent implantation failure (RIF). In these patients, administration of activated PBMC before embryo transfer results in a 2-fold increase in live birth rates(Yang et al., 2020). In this study we evaluated the role of T cells to promotes human endometrial receptivity. On the day of ovulation, PBMC were isolated from and activated with T cells mitogen, the phytohemagglutinin (PHA) and hCG for 48-h in a conditioned culture medium. Distributions of CD4^+^ T cells were characterized in 157 patients by flow cytometry before and after PHA/hCG activation. Cytokine production was analyzed by cytometric beads array. We observed in RIF patients a significant decrease in Th2 and natural Treg cells before activation with PHA/hCG and an increase of Th17 cells after activation compared to intrauterine sperm insemination (IUI) and in vitro fertilization (IVF) groups. Furthermore, the hCG/PHA treatment increases anti-inflammatory T cells (Th2 and Treg cells) compared to non-treated T cells. Principal component analysis (PCA) performed on CD4 T cell subtypes revealed a different cellular profile in the RIF compared to the IUI and IVF groups. This inflammatory state change could explain how endometrium immunomodulation by hCG-activated PBMC helps patients with unexplained RIF to reach implantation.

## 1. Introduction

Nowadays, infertility affects 10 to 15% of couples in industrialized countries (Chandra, Copen, & Stephen, 2014; Evers, 2002). Since the 1980s, we observed a 3-fold increase in the percentage of infertile couples in Canada only (Bushnik, Cook, Yuzpe, Tough, & Collins, 2012). In the last decades, many treatments have been developed in the field of assisted reproductive technology but, based on meta-analysis, only 21 to 30 % of elective single-embryo transfers result in clinical pregnancies (Gleicher, Kushnir, & Barad, 2019)(Gelbaya, Tsoumpou, & Nardo, 2010). A successful pregnancy involves synchronization between a healthy embryo and an endometrial receptivity (Teh, McBain, & Rogers, 2016). Embryo implantation is the critical step leading to the invasion of the uterine wall and the subsequent clinical validation of the pregnancy by increasing human chorionic gonadotropin (hCG) in the maternal blood. Implantation is mediated by interactions between the embryo and the endometrium (Kim & Kim, 2017) and involves hormones, growth factors, cytokines and adhesion molecules (Guzeloglu-Kayisli, Kayisli, & Taylor, 2009; Pawar, Hantak, Bagchi, & Bagchi, 2014). The window of implantation (WOI) occurs between 5 and 10 days post-ovulation and is associated with the expression of cellular products such as cytokines produced by the immune cells (Robertson & Moldenhauer, 2014; van Mourik, Macklon, & Heijnen, 2009). In recent years, studies have shown that a successful embryo implantation maybe influenced by the immune cells present at the implantation site (Robertson, Care, & Moldenhauer, 2018; Robertson & Moldenhauer, 2014).

Recurrent implantation failure (RIF) can be defined as three successive transfers of healthy embryos into a normal uterus that does not lead to successful implantation and clinical pregnancy (Bashiri, Halper, & Orvieto, 2018). RIF can be due to the embryo itself, factors decreasing endometrial receptivity or both (Bashiri et al., 2018). Today, advanced technologies in the field have made it possible to specifically study the implantation process to understand the different aspects of endometrial-embryonic interaction (Hoozemans, Schats, Lambalk, Homburg, & Hompes, 2004; Zohni, Gat, & Librach, 2016). New techniques are being studied to enhance endometrial receptivity. For example, it has been shown that hCG-activated peripheral blood mononuclear cells (PBMC) promote the implantation and invasion of mouse blastocyst *in vitro* (Nakayama et al., 2002). Clinical trials have studied the effects of unstimulated (Okitsu et al., 2011) and hCG-activated PBMC for 48-h on the implantation and clinical pregnancy rates in RIF patients (Yoshioka et al., 2006). These studies have shown that endometrium immunomodulation by intrauterine administration of PBMC significantly increased implantation and clinical pregnancy rates in RIF patients. Furthermore, a recent meta-analysis confirms an increase in live birth rate by 2-fold(Yang et al., 2020). Although PBMC in RIF patients seems to modulate endometrial receptivity, the cellular mechanisms responsible for this phenomenon are still unknown.

Various cell populations such as CD4^+^/CD8^+^ T cells (25-60%), monocytes (10-30%), B cells (5-15%) and NK cells (5-10%) are present in PBMC (Kleiveland, 2015). CD4^+^ T cells are known to play an important role in the establishment of immunity (Luckheeram, Zhou, Verma, & Xia, 2012). These cells perform various functions, characterized by specific cytokine productions that can induce a pro- or anti-inflammatory response. The aim of this study is to understand the role of CD4 T lymphocytes in endometrial immunomodulation in RIF patients.

## 2. Material and Methods

### 2.1 Ethic statement and patients

This study was approved by the ethical committee of INRS (Québec, QC, Canada), protocol CER-16-427. One hundred and fifty-seven (157) women were enrolled in this study between April 2017 and November 2018. All women were enrolled and treated at the Fertilys clinic and are part of a larger cohort. Women were divided into three different groups: 66 patients were treated for intrauterine sperm insemination (**IUI group**), 61 patients with less than three previous embryo implantation failures were transferred with a fresh or frozen/thawed embryo (**IVF group**), and 30 patients with three or more embryo implantation failures were transferred with fresh or frozen/thawed embryos (**RIF group**). Women who had known etiologies of RIF, such as chromosomal abnormalities and the presence of antiphospholipid antibodies were excluded from this study.

### 2.2 In vitro fertilization (IVF) procedure

Ovarian stimulation was performed using mainly an GnRH antagonist protocol combining recombinant FSH/LH to GnRH antagonist (Cetrotide; EMD Serono, Ontario, Canada), and less commonly, a micro-dose flare-up protocol using buserelin as the agonist (Suprefact; Sanofi, Québec, Canada). Briefly, ovarian stimulation was initiated on day 3 using FSH-r (Gonal-F; EMD Serono) or human menopausal gonadotropins (Menopur; Ferring, NJ, USA). For the GnRH antagonist protocol, Cetrotide and rLH (Luveris; EMD Serono) were both added when ovarian follicles reached 12-14 mm in diameter until hCG administration. As for the micro-dose flare-up protocol, the GnRH agonist was started the same day of rFSH or hMG, usually on day 3 of the cycle. When at least two dominant ovarian follicle reached 18 mm in diameter, ovulation was triggered using recombinant or human chorionic gonadotropin (hCG: Ovidrel, EMD Serono or Pregnyl, EMD Serono). Thirty-five to 36-h post-hCG administration, oocytes were recovered by transvaginal oocyte retrieval. Luteal phase was always supported with vaginal progesterone.

### 2.3 Embryo culture

Cumulus-oocyte complexes were removed using hyaluronidase (CooperSurgical Fertility & Genomic Solutions, Målov, Denmark) diluted in modified human tubal fluid (mHTF) media (CooperSurgical Fertility & Genomic Solutions) supplemented with 10% Serum Substitute Supplement (Fujifilm Irvine Scientific, CA, USA). Cumulus-removed oocytes were firstly incubated in Sage 1-Step medium (CooperSurgical Fertility & Genomic Solutions) at 37 °C, 5% CO2, 5% O2. After 2-h, intracytoplasmic sperm injection was performed. Fertilized oocytes were then incubated at 37 °C, 5% CO2, 5% O2 until fresh embryo transfer or embryo vitrification. Depending on woman’s age, one or two embryos were transferred *in utero* three to five days after oocyte retreival.

### 2.4 Frozen/thawed embryo transfer

Embryos were thawed the day before or on the same day of transfer using a vitrification warming kit (Kitazato), according to the manufacturer’s recommendations. Briefly, embryos were placed in a thawing medium for 1 min at 37 °C. After thawing, the embryos were kept in the dilution solution (Kitazato) for 3 min at 37 °C. Embryos were finally washed in the washing solution (Kitazato) for 5 min at 37 °C. Frozen/thawed embryos were transferred *in utero* two days after PBMC intrauterine administration. In most frozen/thawed embryo transfer cycles, letrozole 5 mg from day 3 to 7 was used to augment ovulation followed by hCG administration when the dominant follicle reached 18 mm. Luteal phase was also always supported with vaginal progesterone.

### 2.5 Isolation and administration of PBMC

Thirty milliliter of blood were collected 3 or 5 days prior to embryo transfer and PBMCs were isolated as previously described (Emi et al., 1991; Hashii et al., 1998) by Ficoll-Hypaque (Ficoll-PaqueTM PLUS, GE Healthcare, Uppsala, Sweden). After centrifugation step at 400 g for 30 min, PBMC were washed twice in PBS 1X (Gibco, USA). For immunomodulation, PBMC (1×10^6^ cells/ml) was suspended in 6 ml of global total medium supplemented with PHA/hCG and then incubated for 48-h at 37 °C, 5% CO2. For flow cytometry analysis, 1×10^6^ PBMC of freshly isolated were suspended in 1 ml of RPMI 1640 medium (Gibco) supplemented with 10% fetal bovine serum (FBS), 50 ng/ml Phorbol 12-myristate 13-acetate (PMA), 1 µg/ml ionomycin, 0.7 µl/ml GolgiStop (BD Biosiences, USA) and 1µl/ml GolgiPlug (BD Biosiences) for 5-h at 37 °C, 5% CO2. The remaining of fresh PBMC cells were frozen at -80 °C in FBS supplemented with 10% DMSO. After 2 days, PBMC suspended in global total medium were counted and then washed with new global total medium. The supernatant was frozen at -80 °C. PBMCs (1×10^6^ cells) were suspended in 0.4 ml of global total medium and injected in the uterine cavity. The remaining of PBMCs was suspended in 1 ml of RPMI 1640 (Gibco) supplemented with 10% FBS, 50 ng/ml PMA, 1µg/ml ionomycin, 0.7 µl/ml GolgiStop and 1µl/ml GolgiPlug and were incubated 5-h at 37°C, 5% CO2 for flow cytometry experiments.

### 2.6 Flow Cytometry

After activation of PBMCs with 50 ng/ml PMA, 1 µg/ml ionomycin and 1 µg/ml brefeldin A for 5-h at 37°C, 5% CO2 and a washing step in PBS 1X, 1 x10^6^ cells were transferred into a 1.5 ml microcentrifuge tube. Non-specific binding blocking was achieved by adding 1 µg of normal human IgG (R & D Systems,USA). Staining with anti-CD4^+^/FITC (Biolegend, USA), anti-T-bet/BV605 Biolegend, USA), anti-IFN-γ/APC-Cy7 (Biolegend), anti-GATA-3/BV421 (Biolegend, USA), anti-IL-4/BV510 (Biolegend, USA), anti-IL-17/BV711 (Biolegend, USA), anti-AHR/PE Cy7 (Ebioscience, USA), anti-IL-22/PerCP (Biolegend, USA), anti-CD25/APC (Biolegend, USA), anti-Foxp3/PE (Biolegend, USA). Briefly, cells were stained with cell surface antibodies for 30 min on ice in the dark and then washed with PBS before fixation and permeabilization using Foxp3/Transcription Factor Staining Buffer Set (eBioscience) following the manufacturer’s instructions. Cells were then stained with intracellular antibodies for 30 min at room temperature and washed twice with 1X permeabilization Buffer from the permeabilization kit (eBioscience) before analysis on Fortessa (Becton Dickinson, USA).

### 2.7 Cytometric beads array

Cytokines expression was evaluated using the Human T Helper Cytokine Panel (Biolegend, San Diego, CA) according to the manufacturer’s recommendations. Briefly, the supernatants were diluted with a 1:2 ratio: 25 µl of diluted supernatant were transferred into the 96-well plate, 25 µl of mixed capture beads, 25 µl of detection antibodies cocktail and 25 µl of SA-PE were added into each well. Data were acquired on BD(tm) FACS Fortessa flow cytometer and analyzed with Biolegend legendplex software.

### 2.8 Statistical analysis

Demographic and medical history were compared between the PBMC treated group and the control group using Mann-Whitney tests. Clinical pregnancy and implantation rates were compared between the two groups using chi-squared test. CD4 T cell subsets and cytokines were compared between RIF group and two others groups using Mann-Whitney tests. Statistically significant differences P < 0.05 were considered statistically significant for all tests. CD4 T cell subsets and cytokines were also analysis using PCA model for determining correlations and groups difference for all parameters together. All statistical analysis were performed using GraphPad Prism version 8 (GraphPad Software, San Diego, CA) and RStudio (Open Source Software).

## 3. Results

### 3.1 Stimulation of PHA/hCG of PBMC increases both inflammatory and anti-inflammatory T CD4^+^ subpopulation

In our study, the IUI group, FIV group and RIF group were composed of 66, 61 and 30 women with no significant difference in their age (Table 1). In isolated PBMC, distribution of Th1, Th2, Th17, Th22 and Treg cells populations were evaluated by flow cytometry after 48-h of PHA/hCG stimulation. Analysis of PBMC before *in vitro* stimulation showed that the most abundant CD4^+^ T cells subpopulations were Th1 (11.59%) and Th2 cells (4.49%) (Fig1A and Fig 1B). Following PHA/hCG activation, Th1 cells were slightly, but significantly decreased (−1.04% *p* = 0.041) (Fig. 1A). On the other hand, the proportion of Th2 cells were significantly increased by three times (+10.88%, p<0.0001) following the PHA/hCG exposure (Fig.1B). Th17 cells were also slightly significantly decreased (−0.18%, *p* < 0.0001) in the same condition (Fig.1C). Quite interestingly, the proportion of Th22 cells was significantly increased by 3 times (+2.42%, *p* < 0.0001) (Fig. 1D) and Treg cells, reach a rise of 4 times (+4.07%, *p* < 0.0001) after stimulation (Fig.1E). Thus, PHA/hCG stimulation appears to increased both inflammatory cells (Th22) and anti-inflammatory cells (Th2 and Treg).

**Table 1:**
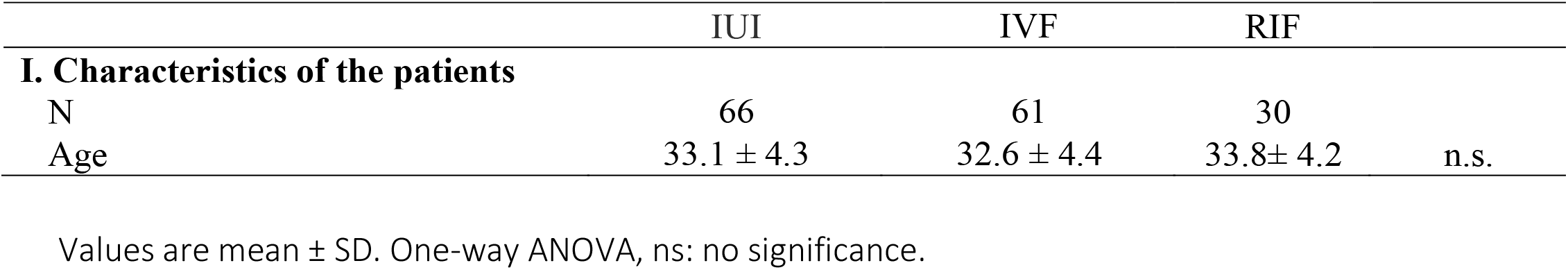
Characteristics of IUI, IVF and RIF patients.

|  | IUI | IVF | RIF |  |
| --- | --- | --- | --- | --- |
| **I. Characteristics of the patients** |  |  |  |  |
| N | 66 | 61 | 30 |  |
| Age | 33.1 ± 4.3 | 32.6 ± 4.4 | 33.8± 4.2 | n.s. |

Values are mean ± SD. One-way ANOVA, ns: no significance.

**Figure 1:**
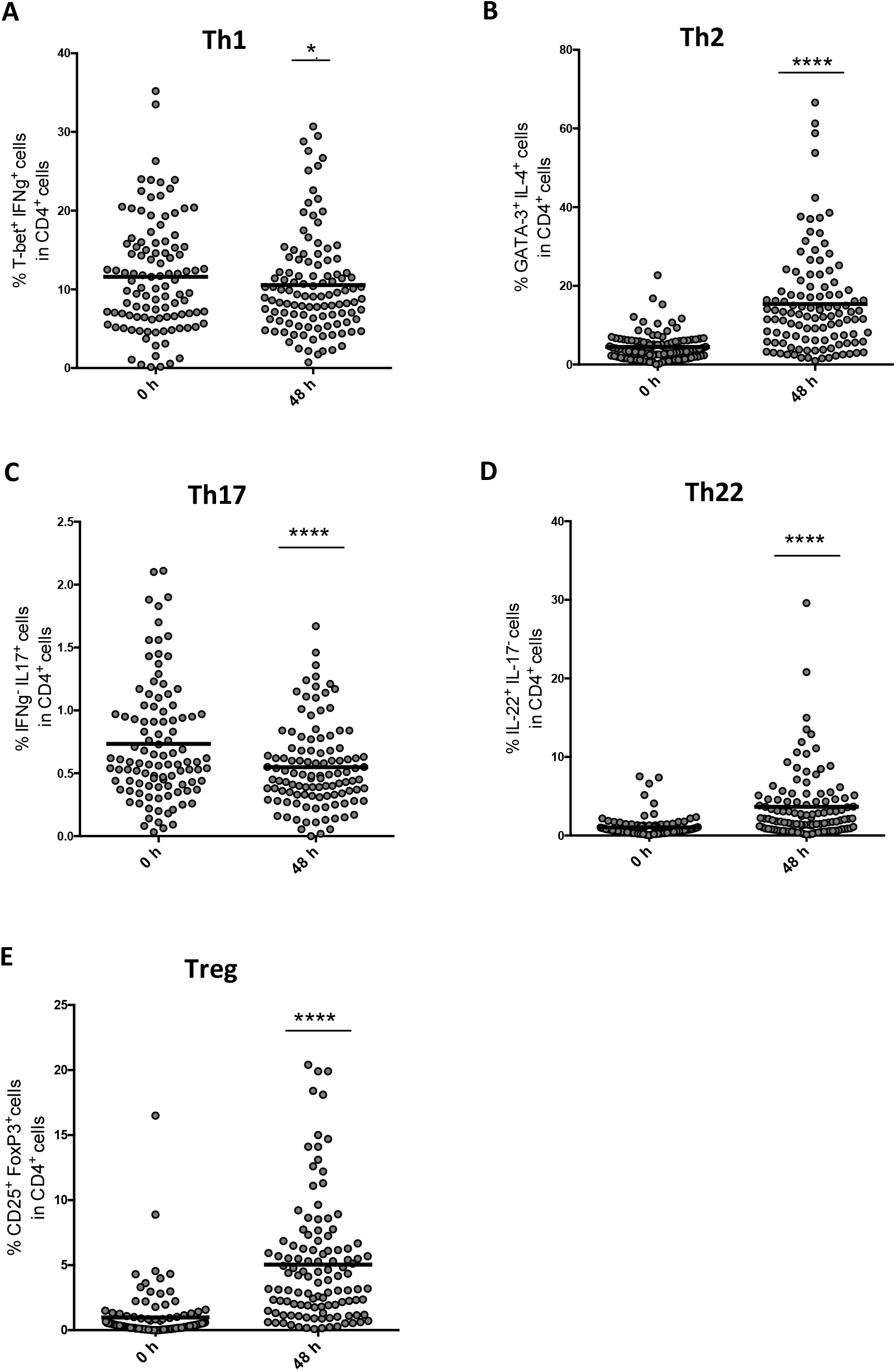
Proportions of lymphocyte subpopulations a) Th1, b) Th2, c) Th17, d) Th22, e) Treg before and after 48h of hCG/PHA activation in our groups of women with reproductive difficulties. Mann-Whitney: p <0.05 (*), p <0.0001 (****), n =157.

### 3.2 Decrease of Th2 cells and natural Treg cells in resting PBMC from RIF patients

In resting PBMC, no significant difference was observed in the distribution of Th1 (Fig. 2A), Th17 (Fig. 2C) and Th22 (Fig. 2D) cells populations between the IUI, IVF and RIF groups of women. In contrast, in RIF population, a decrease of Th2 cells (Fig. 2B, *p* < 0.0001) and Treg cells (Fig. 2E, *p* < 0.001) by around 2,5 and 10 fold respectively, were found when compared to the IUI and IVF group of women.

**Figure 2:**
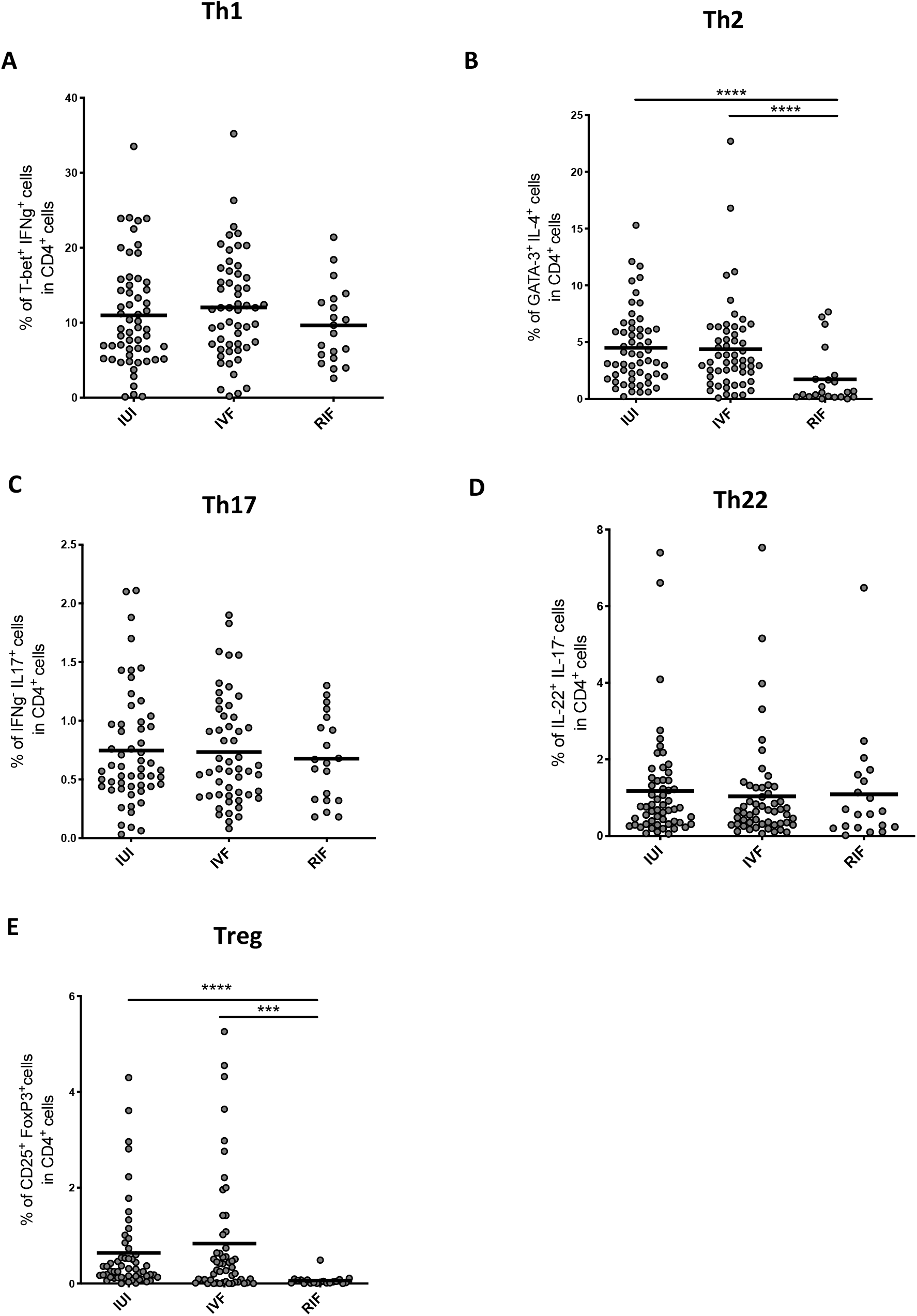
Proportions of lymphocyte subpopulations **(A)** Th1, **(B)** Th2, **(C)** Th17, **(D)** Th22, **(E)** Treg circulant in our patients groups of women with reproductive difficulties. Mann-Whitney: p <0.05 (*), p <0.0001 (****), n = 66 of intrauterine insemination (IUI); n = 61: In vitro fertilization (IVF); n=30: Recurrent implant failure (RIF).

### 3.3 PBMC stimulation by PHA/hCG in RIF patients cause a decrease in Th2 cells and an increase in Th17 cells after treatment

After PHA/hCG activation, no significant difference was observed in the distribution of Th1 (Fig. 3A), Th22 (Fig. 3D), and Treg (Fig. 3E) cells populations between the IUI, IVF and RIF groups of women. In the RIF group, Th2 cells were significantly decreased (*p* < 0.0001) compared to IUI and IVF groups (Fig. 3B). On the other hand, Th17 cells were increased considerably in the RIF group (*p* < 0.01) compared to IUI and IVF groups (Fig. 3C).

**Figure 3:**
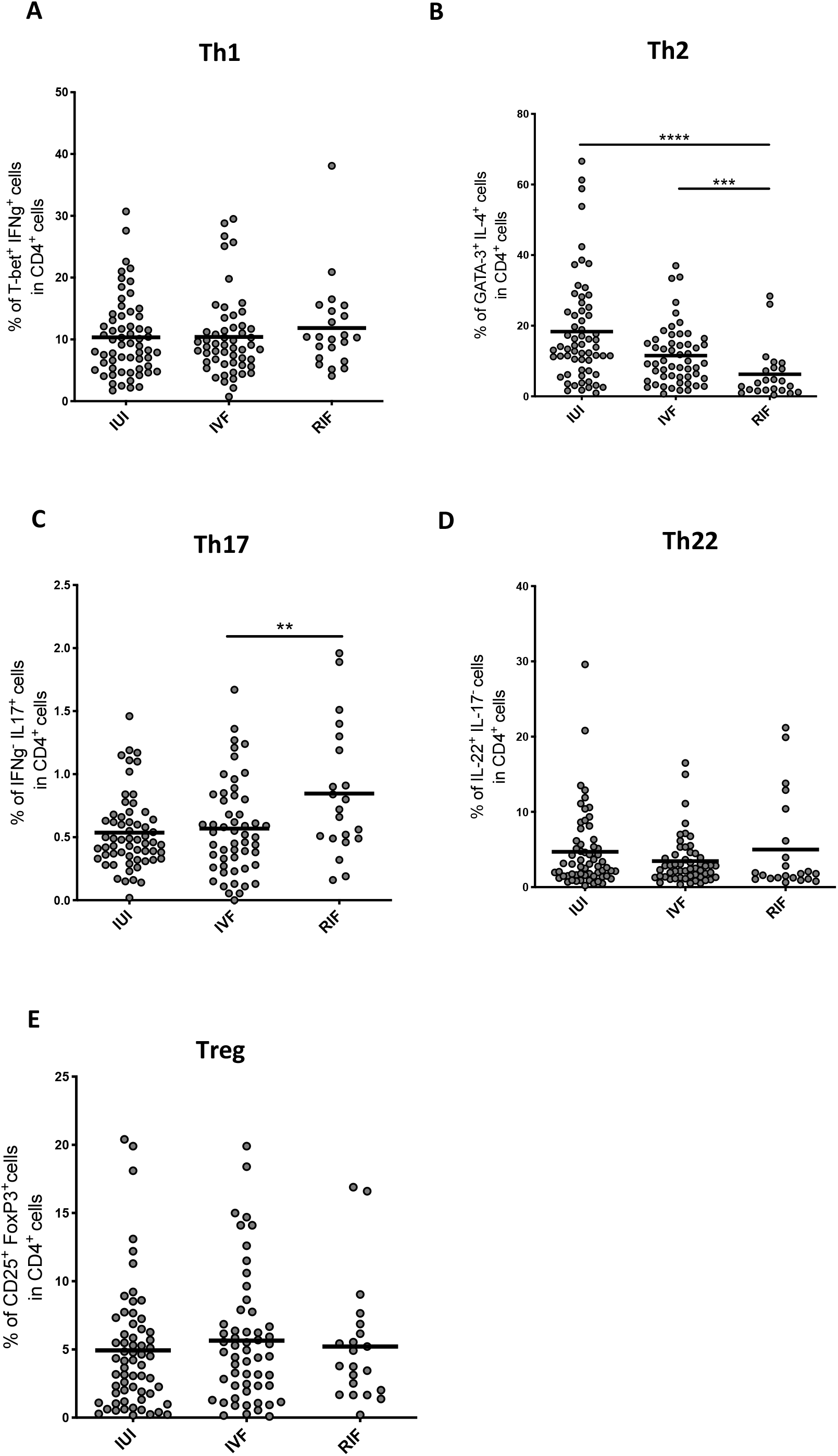
Proportions of lymphocyte subpopulations **(A)** Th1, **(B)** Th2, **(C)** Th17, **(D)** Th22, **(E)** Treg after hCG and PHA treatment in our groups of women with reproductive difficulties. Mann-Whitney: p <0.05 (*), p <0.0001 (****), n = 66 of intrauterine insemination (IUI); n = 61: In vitro fertilization (IVF); n=30: Recurrent implant failure (RIF).

### 3.4 PBMC stimulation by PHA/hCG increases different cytokines productions between RIF, IVF and IUI groups

We analyzed the supernatant of each culture of PHA/hCG stimulated PBMC just before intrauterine administration to determined cytokine profile. For IUI, IVF and RIF groups, PHA/hCG activation led, as expected, to an increase of T cells associated cytokines as compared to unstimulated cells (Fig. 4). PBMC stimulation by PHA/hCG does not affect the production of pro-inflammatory cytokines between the three groups of the patient for IL-2 (Fig. 4A), IFNγ (Fig. 4C), IL17f (Fig. 4E) and IL-22 (Fig. 4K). On the other hand, IL-6 production (Fig. 4B) was increased in the IUI group when compared to IVF (*p* < 0.001) and RIF (*p* < 0.001) groups. For other inflammatory cytokines, IL-17a (Fig. 4D, *p* < 0.05) was increased in the RIF group compared to the IVF group. Interestingly, IL-9 was increased significantly in the RIF group as compared with the other groups (Fig. 4H, *p* < 0.01 for IVF and p< 0.01 for IUI) as was IL-21 in the IVF group compared to IUI and RIF groups (Fig. 4F, *p* < 0.05 for IVF and p< 0.05 for IUI). Regarding anti-inflammatory cytokines, we found only a small decrease of IL-10 in the IVF group compared to the RIF group (Fig. 4K, *p* < 0.05). All other anti-inflammatories cytokines, IL-4 (Fig. 4J), IL-5 (Fig. 4I) and IL-13 (Fig. 4L) were not affected. Thus, globally only IL-6 for the IUI group and IL-9 for the RIF group were significantly increased compared to the other groups.

**Figure 4:**
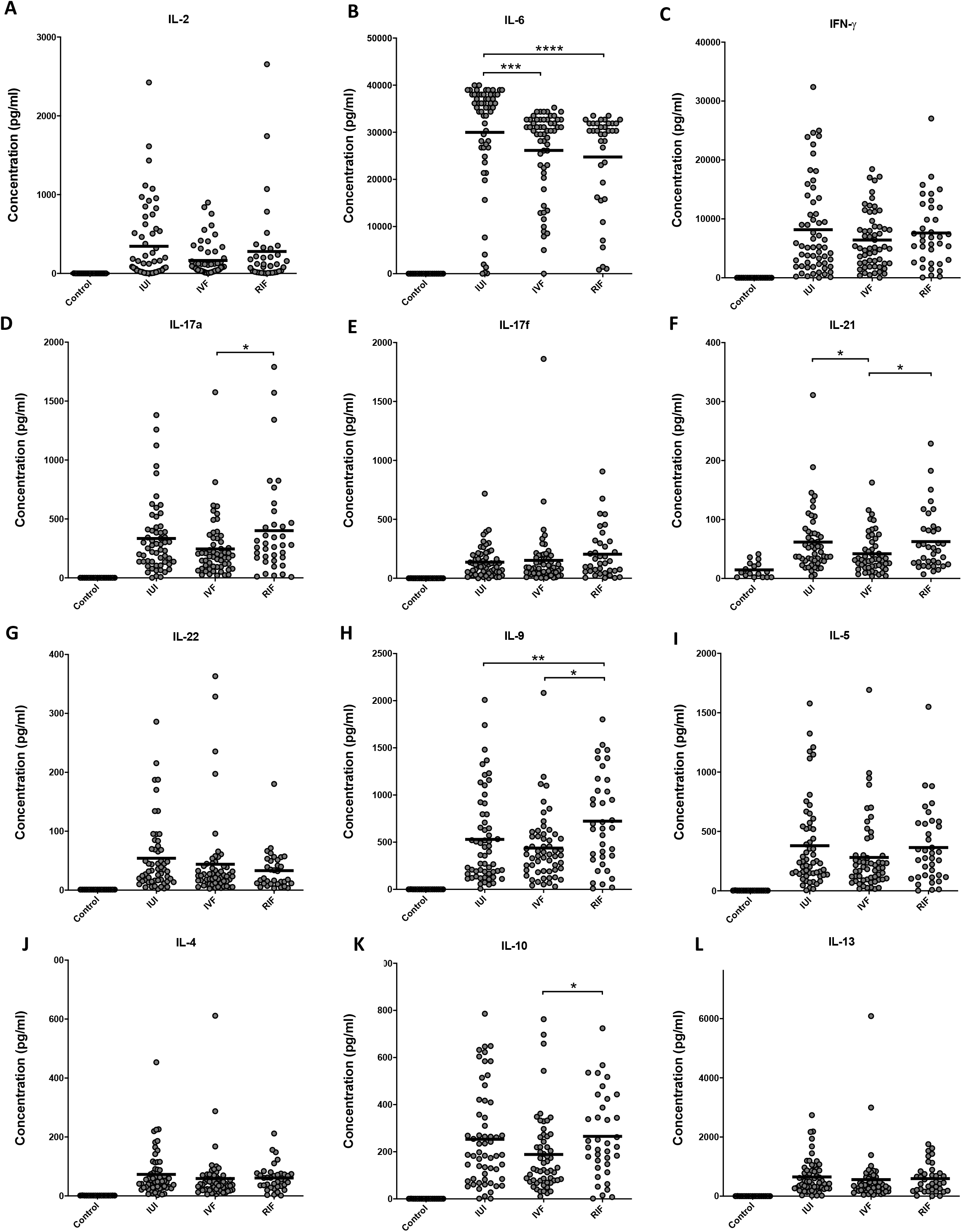
PHA / hCG-treated PBMC induces an increase of CD4 T cells cytokines. The amount of **(A)** IL-2 secreted, **(B)** IL-6 secreted, **(C)** IFN-γ secreted, **(D)** IL-17a secreted, **(E)** IL-17f secreted, **(F)** IL-21 secreted, **(G)** IL-22 secreted, **(H)** IL-9 secreted **(I)** IL-5secreted, **(J)** IL-4 secreted, **(K)** IL-10 secreted **(L)** IL-13 secreted by PBMC treated with PHA / hCG for 48 hours in IUI group, IVF group and RIF group. Mann-Whitney: p <0.05 (*), p <0.0001 (****), n = 66 for Intrauterine Insemination (IUI); n = 61 for In vitro Fertilization (IVF); n=30 for Recurrent Implantation Failure (RIF), n=17 for Control group.

### 3.5 Differences between RIF patients and other patients shown by PCA on all the data

To obtain a better analysis of our results, we used principal component analysis (PCA) on all of our data. The PCA method allows treating several potentially linked variables into a new variable defined as the linear component of the original variables. When we integrated our cell subpopulations data for unstimulated cells recovery at day 0, we see that the patient groups IUI and IVF were similar, their point clouds were located in the same regions of the model (Fig 5A). In contrast, the RIF group of women has a different representation of their variables. By using the data of the CD4^+^ T cell subpopulations after PHA/hCG stimulation, as well as the cytokines produced by the cells in the PCA model, we obtained a reduction in the differences observed between our patients (Fig 5B). When we performed a spearman correlation on all the data of T-cell subpopulations and cytokines analyzed, we observed several strong positive correlations among the anti-inflammatory cytokines of the Th2 phenotype (IL-4, IL-5, IL-9 and IL-13) (Fig 5C and 5D). Moreover, there was a positive correlation between the cytokines IL-17a, IL-17f and IL-22, associated with the Th17 phenotype (Fig 5C and 5D). We do not observed correlations between the T cell subpopulations after PHA/hCG stimulation Fig 5D.

**Figure 5:**
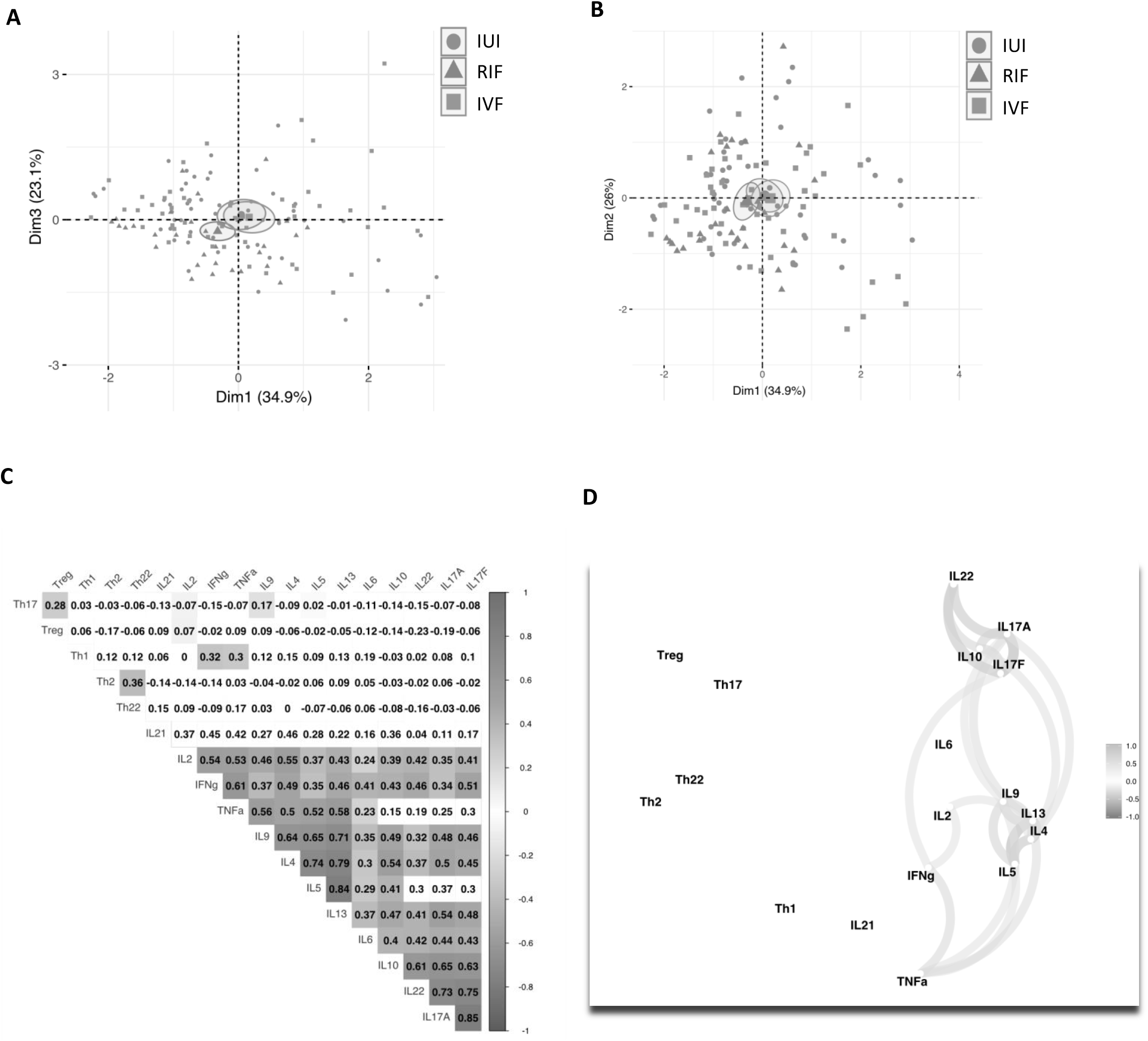
Plot of individuals of the multiple factor analysis (MFA) applied on T cells subsets at **(A)** before hCG/PHA treatment and **(B)** after hCG/PHA treatment. IUI patients, RIF patients and IVF patients are represented respectively by the red, green and blue dots. **(C)** Spearman correlation coefficients of T cells subsets and T cells cytokines at J2. Representation of correlation clusters of T cells subsets and T cells cytokines at day 2.

### 3.6 Administration of activated PBMC by PHA/hCG does not affect IUI and IVF groups

To evaluate the impact of activated *PBMC by PHA/hCG* in improving embryo implantation,we have compared IUI and IVF groups. Age, number of previous embryo transfer(s), number of embryos transferred, and previous IUI were not statistically different between the PBMC-treated and control groups (Table 1S and Table 2S). Implantation and clinical pregnancy rates were not statistically significant either between the PBMC treated vs non-treated in IUI and IVF patients’ groups.

## 4. Discussion

In this study, we showed in PBMC a decrease of anti-inflammatory cells (Th2, Treg) in the RIF group. Stimulation of PBMC with PHA/hCG in the RIF group causes a slight rise of Th2 and normalization of Treg cells. Associated with later change, % Th17 were also predominant in the RIF group. Concerning cytokine produce by PBMC stimulated by PHA/hCG, only IL-9 was different between RIF patients and IUI/IVF patients. The embryo implantation is one of the significant steps in the success of a pregnancy. Although, with the progress made in embryo selection methods and the detection of possible mutations, which may disadvantage the implantation process, the endometrial receptivity remains a problematic element to assess (Katzorke, Vilella, Ruiz, Krussel, & Simon, 2016). To explain the role of PBMCs in the implantation process, several possible mechanisms have been suggested. Progesterone is secreted during the luteal phase to prepare the endometrium for implantation of the embryo (Mesen & Young, 2015). It has been suggested that PBMCs administered into the uterine cavity move to the endometrial stromal tissue, promoting attachment and embryo invasion (Hiroshi Fujiwara et al., 2009). PBMCs have been shown to stimulate the production of progesterone, in particular, via the action of Th2 cells (Hashii et al., 1998). Our results show that the hCG/PHA treatment promotes Th2 and Treg cells, associated with an anti-inflammatory response. The presence of anti-inflammatory cells is necessary to reduce inflammation after embryo implantation (Sykes, MacIntyre, Yap, Teoh, & Bennett, 2012; Tsuda, Nakashima, Shima, & Saito, 2019). The production of leukemia inhibitory factor and endometrial vascular endothelial growth factor is increased by PBMC used for immunomodulation (Yu et al., 2014). It also appears that PBMC treatment increases the invasion capacity of the trophoblast cells and the differentiation of corpus luteum and endometrium (Campesato et al., 2020).

The diagnosis of endometrial receptivity represents a significant challenge. Several tests have been suggested to overcome this challenge. MicroArray technology has been proposed to evaluate the transcriptomic signature of the window of implantation (WOI) (Herington, Guo, Reese, & Paria, 2016; Schmidt et al., 2005). It was initially developed as a diagnostic tool for endoplasmic reticulum stress (Hamamura, Liu, & Yokota, 2008). This technology has made it possible to demonstrate that the WOI was displaced in 25% of RIF patients (Ruiz-Alonso et al., 2013). A personalized embryo transfer to improve reproductive performance can be obtained with this method (Campesato et al., 2020). Although this technology increases the implantation rate in some patients due to a better understanding of the WOI, other factors have a considerable impact on the success of the embryo implantation and its clinical usefulness remains to be proven.

The data in the literature indicates that a pro-inflammatory environment is often found in women who have had RIF (Liang et al., 2015). This pro-inflammatory state present in the endometrium may cause a decline in endometrium receptivity and thus explain the inability of the embryo to implant. Besides, a study showed that the presence of Treg cells increases pregnancy rates in RIF patients (Ahmadi et al., 2017). Our results showed that Treg cells were significantly reduced in the RIF group. In addition, Th2 cells, also anti-inflammatory, are strongly reduced in the RIF group. An anti-inflammatory state is essential for the maintenance of the embryo after implantation (Sykes et al., 2012; Tsuda et al., 2019), and a decrease in these cells may explain the reduction in receptivity in the RIF group (Ahmadi et al., 2017).

Moreover, hCG/PHA treatment increases induced Treg cells differentiation to make it similar to patients in IUI and IVF groups. Induced Tregs have a regulatory capacity similar to natural Treg cells (Shevach & Thornton, 2014). The treatment therefore promotes anti-inflammatory cell differentiation, helping to reduce post-implantation inflammation and maintain embryo growth.

Pro-inflammatory and anti-inflammatory responses both play a role in the success of pregnancy. A study has shown that in the plasma of RIF patients, a large quantity of pro-inflammatory cytokines (IFN-γ, IL-6, IL-1β), as well as a decrease in TGfβ1 can be found (Campesato et al., 2020). In our study we did not observe differences between RIF patients and IUI and IVF groups in the differentiation of Th17 cells, whose cytokines IL-6 and IL-1β promote differentiation and Th1 cells, wich produce IFN-γ. On the other hand, we observed a substantial decrease in Treg cells in RIF patients than the other groups. This decrease could be explained by the TGFβ levels found in RIF patients.

In most immunomodulation protocols for RIF patients’ treatment, PBMC cells used are either unstimulated or stimulated with hCG (Okitsu et al., 2011; Yoshioka et al., 2006). The quantity of cells used ranges from 1 to 2 x10^7^ cells (Yoshioka et al., 2006). Our approach was to use PBMC cells activated by PHA and hCG treatment with a lower number of cells, of the order of 1×10^6^ cells. We observed that activating PBMC by this protocol not only induced the T cell subpopulation associated with an anti-inflammatory ativity but without effect on pro-inflammatory cells.

Immunomodulation using PBMCs has been shown to increase the live birth rate in RIF patients (Yoshioka et al., 2006)(Yang et al., 2020). On the other hand, in patients considered non-RIF, this treatment does not seem to increase the pregnancy success rate (Li et al., 2017; Okitsu et al., 2011). In the present study, we observed that there was also no increase in clinical pregnancy rate in IUI and IVF groups with PBMC treatment (Table 1S and 2S). Sample sizes of our studied populations are too small to draw any conclusions in that regard.

An increase of IL-9 in the supernatant of PHA/hCG-stimulated PBMC in the RIF group suggests a potential role of this interleukin in successful implantation. The primary source of IL-9 is Th9 lymphocytes, which is differentiated in the presence of TGFβ and IL-4 (Goswami & Kaplan, 2011). Although the Th9 subpopulation was not analyzed in our study, their increase seems logical since we observe a rise in TGFβ (Treg) and IL-4 (Th2) producing cells after PHA/hCG stimulation. Moreover, Th9 and Th2 cells have a similar requirement for several transcription factors, including STAT6, GATA3 and IRF4 (Jabeen et al., 2013). IL-9 is a major cytokine at the mouse fetal-maternal interface (Habbeddine, Verbeke, Karaz, Bobe, & Kanellopoulos-Langevin, 2014) and plays a vital role in regulating local inflammatory immune responses and promoting allograft tolerance (Wan et al., 2020; Wang, Sung, Gilman-Sachs, & Kwak-Kim, 2020). Another possible function of IL-9 in the implantation process can be associated with their capacity to induce the stimulation of CCL20 production (Habbeddine et al., 2014). In pig model, it was demonstrated that activation of CCL20 and its receptor CCR6 promotes endometrium preparation for implantation (Park, Bae, Bazer, Song, & Lim, 2019). Another important role of CCL20 is their involvement in the recruitment of decidual immune cell subtypes such as dendritic cells and Treg from peripheral tissues (Du, Wang, & Li, 2014).

Depending on the studied populations (IUI, IVF or RIF), this study have demonstrated significant differences in PBMC profiles, before and after in vitro activation, both at the levels of cell differentiation and cytokine production, that could help to understand how endometrium immunomodulation by hCG-activated PBMC helps patients with unexplained RIF to reach implantation. The immunomodulation of the endometrium by PBMCs can be used as an immunological therapeutic approach in addition to the other therapies already used. (H. Fujiwara et al., 2016). Also, the use of PBMC in these treatments should not lead to an over activation of the uterine immune profile at the risk of becoming harmful for the success of pregnancy. (Ledee et al., 2017). This is why the use of a cell stimulation protocol promoting an anti-inflammatory profile is necessary when the patient has a pro-inflammatory profile. Determining the uterine inflammatory profile would make it possible to apply treatments that are appropriate to the needs of each patient.

## Acknowledgments

The authors are grateful to the Fertilys embryologists and nurses for their implication in this project. Mer Kulbay for the edition.

## References

Ahmadi, M., Abdolmohammadi-Vahid, S., Ghaebi, M., Aghebati-Maleki, L., Dolati, S., Farzadi, L., … Yousefi, M. (2017). Regulatory T cells improve pregnancy rate in RIF patients after additional IVIG treatment. Syst Biol Reprod Med, 63(6), 350–359. doi:10.1080/19396368.2017.1390007

Bashiri, A., Halper, K. I., & Orvieto, R. (2018). Recurrent Implantation Failure-update overview on etiology, diagnosis, treatment and future directions. Reprod Biol Endocrinol, 16(1), 121. doi:10.1186/s12958-018-0414-2

Bushnik, T., Cook, J. L., Yuzpe, A. A., Tough, S., & Collins, J. (2012). Estimating the prevalence of infertility in Canada. Hum Reprod, 27(3), 738–746. doi:10.1093/humrep/der465

Campesato, L. F., Budhu, S., Tchaicha, J., Weng, C. H., Gigoux, M., Cohen, I. J., … Wolchok, J.D. (2020). Blockade of the AHR restricts a Treg-macrophage suppressive axis induced by L-Kynurenine. Nat Commun, 11(1), 4011. doi:10.1038/s41467-020-17750-z

Chandra, A., Copen, C. E., & Stephen, E. H. (2014). Infertility service use in the United States: data from the National Survey of Family Growth, 1982-2010. Natl Health Stat Report(73), 1–21. Retrieved from https://www.ncbi.nlm.nih.gov/pubmed/24467919

Du, M. R., Wang, S. C., & Li, D. J. (2014). The integrative roles of chemokines at the maternal-fetal interface in early pregnancy. Cell Mol Immunol, 11(5), 438–448. doi:10.1038/cmi.2014.68

Emi, N., Kanzaki, H., Yoshida, M., Takakura, K., Kariya, M., Okamoto, N., … Mori, T. (1991). Lymphocytes stimulate progesterone production by cultured human granulosa luteal cells. Am J Obstet Gynecol, 165(5 Pt 1), 1469–1474. doi:10.1016/0002-9378(91)90393-6

Evers, J. L. (2002). Female subfertility. Lancet, 360(9327), 151–159. doi:10.1016/S0140-6736(02)09417-5

Fujiwara, H., Araki, Y., Imakawa, K., Saito, S., Daikoku, T., Shigeta, M., … Mori, T. (2016). Dual Positive Regulation of Embryo Implantation by Endocrine and Immune Systems--Step-by-Step Maternal Recognition of the Developing Embryo. Am J Reprod Immunol, 75(3), 281–289. doi:10.1111/aji.12478

Fujiwara, H., Ideta, A., Araki, Y., Takao, Y., Sato, Y., Tsunoda, N., … Konishi, I. (2009). Immune System Cooperatively Supports Endocrine System-Primed Embryo Implantation. Journal of Mammalian Ova Research, 26(3), 122–128. doi:10.1274/jmor.26.122

Gelbaya, T. A., Tsoumpou, I., & Nardo, L. G. (2010). The likelihood of live birth and multiple birth after single versus double embryo transfer at the cleavage stage: a systematic review and meta-analysis. Fertil Steril, 94(3), 936–945. doi:10.1016/j.fertnstert.2009.04.003

Gleicher, N., Kushnir, V. A., & Barad, D. H. (2019). Worldwide decline of IVF birth rates and its probable causes. Hum Reprod Open, 2019(3), hoz017. doi:10.1093/hropen/hoz017

Goswami, R., & Kaplan, M. H. (2011). A brief history of IL-9. J Immunol, 186(6), 3283–3288. doi:10.4049/jimmunol.1003049

Guzeloglu-Kayisli, O., Kayisli, U. A., & Taylor, H. S. (2009). The role of growth factors and cytokines during implantation: endocrine and paracrine interactions. Semin Reprod Med, 27(1), 62–79. doi:10.1055/s-0028-1108011

Habbeddine, M., Verbeke, P., Karaz, S., Bobe, P., & Kanellopoulos-Langevin, C. (2014). Leukocyte population dynamics and detection of IL-9 as a major cytokine at the mouse fetal-maternal interface. PLoS One, 9(9), e107267. doi:10.1371/journal.pone.0107267

Hamamura, K., Liu, Y., & Yokota, H. (2008). Microarray analysis of thapsigargin-induced stress to the endoplasmic reticulum of mouse osteoblasts. J Bone Miner Metab, 26(3), 231–240. doi:10.1007/s00774-007-0825-1

Hashii, K., Fujiwara, H., Yoshioka, S., Kataoka, N., Yamada, S., Hirano, T., … Maeda, M. (1998). Peripheral blood mononuclear cells stimulate progesterone production by luteal cells derived from pregnant and non-pregnant women: possible involvement of interleukin-4 and interleukin-10 in corpus luteum function and differentiation. Hum Reprod, 13(1O), 2738–2744. doi:10.1093/humrep/13.10.2738

Herington, J. L., Guo, Y., Reese, J., & Paria, B. C. (2016). Gene profiling the window of implantation: Microarray analyses from human and rodent models. J Reprod Health Med, 2(Suppl 2), S19–S25. doi:10.1016/j.jrhm.2016.11.006

Hoozemans, D. A., Schats, R., Lambalk, C. B., Homburg, R., & Hompes, P. G. (2004). Human embryo implantation: current knowledge and clinical implications in assisted reproductive technology. Reprod Biomed Online, 9(6), 692–715. doi:10.1016/s1472-6483(10)61781-6

Jabeen, R., Goswami, R., Awe, O., Kulkarni, A., Nguyen, E. T., Attenasio, A., … Kaplan, M. H. (2013). Th9 cell development requires a BATF-regulated transcriptional network. J Clin Invest, 123(11), 4641–4653. doi:10.1172/JCI69489

Katzorke, N., Vilella, F., Ruiz, M., Krussel, J. S., & Simon, C. (2016). Diagnosis of Endometrial-Factor Infertility: Current Approaches and New Avenues for Research. Geburtshilfe Frauenheilkd, 76(6), 699–703. doi:10.1055/s-0042-103752

Kim, S. M., & Kim, J. S. (2017). A Review of Mechanisms of Implantation. Dev Reprod, 21(4), 351–359. doi:10.12717/DR.2017.21.4.351

Kleiveland, C. R. (2015). Peripheral Blood Mononuclear Cells. In K. Verhoeckx, P. Cotter, I. Lopez-Exposito, C. Kleiveland, T. Lea, A. Mackie, T. Requena, D. Swiatecka, & H. Wichers (Eds.), The Impact of Food Bioactives on Health: in vitro and ex vivo models (pp. 161–167). Cham (CH).

Ledee, N., Prat-Ellenberg, L., Chevrier, L., Balet, R., Simon, C., Lenoble, C., … Petitbarat, M. (2017). Uterine immune profiling for increasing live birth rate: A one-to-one matched cohort study. J Reprod Immunol, 119, 23–30. doi:10.1016/j.jri.2016.11.007

Li, S., Wang, J., Cheng, Y., Zhou, D., Yin, T., Xu, W., … Yang, J. (2017). Intrauterine administration of hCG-activated autologous human peripheral blood mononuclear cells (PBMC) promotes live birth rates in frozen/thawed embryo transfer cycles of patients with repeated implantation failure. J Reprod Immunol, 119, 15–22. doi:10.1016/j.jri.2016.11.006

Liang, P. Y., Diao, L. H., Huang, C. Y., Lian, R. C., Chen, X., Li, G. G., … Zeng, Y. (2015). The pro-inflammatory and anti-inflammatory cytokine profile in peripheral blood of women with recurrent implantation failure. Reprod Biomed Online, 31(6), 823–826. doi:10.1016/j.rbmo.2015.08.009

Luckheeram, R. V., Zhou, R., Verma, A. D., & Xia, B. (2012). CD4(+)T cells: differentiation and functions. Clin Dev Immunol, 2012, 925135. doi:10.1155/2012/925135

Mesen, T. B., & Young, S. L. (2015). Progesterone and the luteal phase: a requisite to reproduction. Obstet Gynecol Clin North Am, 42(1), 135–151. doi:10.1016/j.ogc.2014.10.003

Nakayama, T., Fujiwara, H., Maeda, M., Inoue, T., Yoshioka, S., Mori, T., & Fujii, S. (2002). Human peripheral blood mononuclear cells (PBMC) in early pregnancy promote embryo invasion in vitro: HCG enhances the effects of PBMC. Human Reproduction, 17(1), 207–212. doi:DOI 10.1093/humrep/17.1.207

Okitsu, O., Kiyokawa, M., Oda, T., Miyake, K., Sato, Y., & Fujiwara, H. (2011). Intrauterine administration of autologous peripheral blood mononuclear cells increases clinical pregnancy rates in frozen/thawed embryo transfer cycles of patients with repeated implantation failure. J Reprod Immunol, 92(1-2), 82–87. doi:10.1016/j.jri.2011.07.001

Park, C., Bae, H., Bazer, F. W., Song, G., & Lim, W. (2019). Activation of CCL20 and its receptor CCR6 promotes endometrium preparation for implantation and placenta development during the early pregnancy period in pigs. Dev Comp Immunol, 92, 35–42. doi:10.1016/j.dci.2018.11.005

Pawar, S., Hantak, A. M., Bagchi, I. C., & Bagchi, M. K. (2014). Minireview: Steroid-regulated paracrine mechanisms controlling implantation. Mol Endocrinol, 28(9), 1408–1422. doi:10.1210/me.2014-1074

Robertson, S. A., Care, A. S., & Moldenhauer, L. M. (2018). Regulatory T cells in embryo implantation and the immune response to pregnancy. J Clin Invest, 128(10), 4224–4235. doi:10.1172/JCI122182

Robertson, S. A., & Moldenhauer, L. M. (2014). Immunological determinants of implantation success. Int J Dev Biol, 58(2-4), 205–217. doi:10.1387/ijdb.140096sr

Ruiz-Alonso, M., Blesa, D., Diaz-Gimeno, P., Gomez, E., Fernandez-Sanchez, M., Carranza, F., … Simon, C. (2013). The endometrial receptivity array for diagnosis and personalized embryo transfer as a treatment for patients with repeated implantation failure. Fertil Steril, 100(3), 818–824. doi:10.1016/j.fertnstert.2013.05.004

Schmidt, A., Groth, P., Haendler, B., Hess-Stumpp, H., Krätzschmar, J., Seidel, H., … Weiss, B. (2005). Gene Expression During the Implantation Window: Microarray Analysis of Human Endometrial Samples, Berlin, Heidelberg.

Shevach, E. M., & Thornton, A. M. (2014). tTregs, pTregs, and iTregs: similarities and differences. Immunol Rev, 259(1), 88–102. doi:10.1111/imr.12160

Sykes, L., MacIntyre, D. A., Yap, X. J., Teoh, T. G., & Bennett, P. R. (2012). The Th1:th2 dichotomy of pregnancy and preterm labour. Mediators Inflamm, 2012, 967629. doi:10.1155/2012/967629

Teh, W. T., McBain, J., & Rogers, P. (2016). What is the contribution of embryo-endometrial asynchrony to implantation failure? J Assist Reprod Genet, 33(11), 1419–1430. doi:10.1007/s10815-016-0773-6

Tsuda, S., Nakashima, A., Shima, T., & Saito, S. (2019). New Paradigm in the Role of Regulatory T Cells During Pregnancy. Front Immunol, 10, 573. doi:10.3389/fimmu.2019.00573

van Mourik, M. S., Macklon, N. S., & Heijnen, C. J. (2009). Embryonic implantation: cytokines, adhesion molecules, and immune cells in establishing an implantation environment. J Leukoc Biol, 85(1), 4–19. doi:10.1189/jlb.0708395

Wan, J., Wu, Y., Ji, X., Huang, L., Cai, W., Su, Z., … Xu, H. (2020). IL-9 and IL-9-producing cells in tumor immunity. Cell Commun Signal, 18(1), 50. doi:10.1186/s12964-020-00538-5

Wang, W., Sung, N., Gilman-Sachs, A., & Kwak-Kim, J. (2020). T Helper (Th) Cell Profiles in Pregnancy and Recurrent Pregnancy Losses: Th1/Th2/Th9/Th17/Th22/Tfh Cells. Front Immunol, 11, 2025. doi:10.3389/fimmu.2020.02025

Yang, D. N., Wu, J. H., Geng, L., Cao, L. J., Zhang, Q. J., Luo, J. Q., … Xia, X. (2020). Efficacy of intrauterine perfusion of peripheral blood mononuclear cells (PBMC) for infertile women before embryo transfer: meta-analysis. J Obstet Gynaecol, 40(7), 961–968. doi:10.1080/01443615.2019.1673711

Yoshioka, S., Fujiwara, H., Nakayama, T., Kosaka, K., Mori, T., & Fujii, S. (2006). Intrauterine administration of autologous peripheral blood mononuclear cells promotes implantation rates in patients with repeated failure of IVF–embryo transfer. Human Reproduction, 21(12), 3290–3294. doi:10.1093/humrep/del312

Yu, N., Yang, J., Guo, Y., Fang, J., Yin, T., Luo, J., … Xu, W. (2014). Intrauterine administration of peripheral blood mononuclear cells (PBMCs) improves endometrial receptivity in mice with embryonic implantation dysfunction. Am J Reprod Immunol, 71(1), 24–33. doi:10.1111/aji.12150

Zohni, K. M., Gat, I., & Librach, C. (2016). Recurrent implantation failure: a comprehensive review. Minerva Ginecol, 68(6), 653-667. Retrieved from https://www.ncbi.nlm.nih.gov/pubmed/26982235

