## SUPPLEMENTARY DATA for "Role of T cells in intrauterine administration of activated peripheral blood mononuclear cells in recurrent implantation failure" (from another version of this preprint: version 1, 2021-01-08)

**Supplemental table 1: Characteristics and clinical outcomes of IUI patients treated or non-treated with peripheral blood mononuclear cells (PBMC).** Characteristics analysis was performed using Man-Whitney. Clinical outcome analysis was performed using Chi-square. Values are mean ± SD. ns: no significance.

| | PBMC-treated | Non-treated | |
|---|---|---|---|
| **I. Characteristics** | | | |
| N | 30 | 36 | |
| Age | 33.3 ± 4.4 | 32.8 ± 4.2 | n.s. |
| No. of previous IUI attempts | 1.78 ± 1.37 | 2.3 ± 1.46 | n.s. |
| **II. Clinical outcome** | | | |
| Clinical pregnancy rate | 13.3% (4/30) | 5.6% (2/36) | n.s. |
| Implantation rate | 6.7% (2/30) | 2.8% (1/36) | n.s. |

**Supplemental table 2: Characteristics and clinical outcomes of IVF patients treated or non-treated with peripheral blood mononuclear cells (PBMC).** Characteristics analysis was performed using Man-Whitney. Clinical outcome analysis was performed using Chi-square. Values are mean ± SD ns: no significance.

| | PBMC treated | Non-treated | |
|---|---|---|---|
| **I. Characteristics** | | | |
| N | 28 | 33 | |
| Age | 33.3 ± 3.6 | 32,1 ± 5.0 | n.s. |
| IVF–embryo transfer previous attempts | 0.57 ± 0.74 | 0.68 ± 0.74 | n.s. |
| No. of embryo transferred | 1.07 ± 0.262 | 1.03 ± 0.17 | n.s. |
| **II. Clinical outcome** | | | |
| Clinical pregnancy rate | 35.7% (10/28) | 51.5% (17/33) | n.s. |
| Implantation rate | 32.1% (9/28) | 51.5% (17/33) | n.s. |

**Supplemental table 3: Characteristics and clinical outcomes of RIF patients in the peripheral blood mononuclear cells (PBMC)-treated group.** Values are mean ± SD.

| | PBMC treated |
|---|---|
| **I. Characteristics** | |
| N | 30 |
| Age | 33.8 ± 4.2 |
| IVF–embryo transfer previous attempts | 3.93 ± 2.71 |
| No. of embryo transferred | 1.07 ± 0.254 |
| **II. Clinical outcome** | |
| Clinical pregnancy rate | 23.3 % (7/30) |
| Implantation rate | 23.3 % (7/30) |
